# A homozygous variant in mitochondrial RNase P subunit PRORP is associated with Perrault syndrome characterized by hearing loss and primary ovarian insufficiency

**DOI:** 10.1101/168252

**Authors:** Irit Hochberg, Leigh A. M. Demain, Jill E. Urquhart, Albert Amberger, Andrea J. Deutschmann, Sandra Demetz, Kyle Thompson, James O'sullivan, Inna A. Belyantseva, Melanie Barzik, Simon G. Williams, Sanjeev S. Bhaskar, Emma M. Jenkinson, Nada AlSheqaih, Zeev Blumenfeld, Sergey Yalonetsky, Stephanie Oerum, Walter Rossmanith, Wyatt W. Yue, Johannes Zschocke, Robert W. Taylor, Thomas B. Friedman, Kevin J. Munro, Raymond T. O'Keefe, William G. Newman

## Abstract

Perrault syndrome is a rare autosomal recessive condition characterised by sensorineural hearing loss in both sexes and primary ovarian insufficiency in 46 XX, females. It is genetically heterogeneous with biallelic variants in six genes identified to date (*HSD17B4*, *HARS2*, *LARS2*, *CLPP*, *C10orf2* and *ERAL1*). Most genes possessing variants associated with Perrault syndrome are involved in mitochondrial translation. We describe a consanguineous family with three affected individuals homozygous for a novel missense variant c.1454C>T; p.(Ala485Val) in *KIAA0391*, encoding proteinaceous RNase P (PRORP), the metallonuclease subunit of the mitochondrial RNase P complex, responsible for the 5’-end processing of mitochondrial precursor tRNAs. In RNase P activity assays, RNase P complexes containing the PRORP disease variant produced ~35-45% less 5’-processed tRNA than wild type PRORP. Consistently, the accumulation of unprocessed polycistronic mitochondrial transcripts was observed in patient dermal fibroblasts, leading to an observable loss of steady-state levels of mitochondrial oxidative phosphorylation components. Expression of wild type *KIAA0391* in patient fibroblasts rescued tRNA processing. Immunohistochemistry analyses of the auditory sensory epithelium from postnatal and adult mouse inner ear showed a high level of PRORP in the efferent synapses and nerve fibres of hair cells, indicating a possible mechanism for the sensorineural hearing loss observed in affected individuals. We have identified a variant in an additional gene associated with Perrault syndrome. With the identification of this disease-causing variant in *KIAA0391*, reduced function of each of the three subunits of mitochondrial RNase P have now been associated with distinct clinical presentations.

**Author Summary:** Perrault syndrome is a rare genetic condition which results in hearing loss in both sexes and ovarian dysfunction in females. Perrault syndrome may also cause neurological symptoms in some patients. Here, we present the features and genetic basis of the condition in three sisters affected by Perrault syndrome. The sisters did not have pathogenic variants in any of the genes previously associated with Perrault syndrome. We identified a change in the gene *KIAA0391*, encoding PRORP, a subunit of the mitochondrial RNase P complex. Mitochondrial RNase P is a key enzyme in RNA processing in mitochondria. Impaired RNA processing reduces protein production in mitochondria, which we observed in patient cells along with high levels of unprocessed RNA. When we expressed wild type PRORP in patient cells, the RNA processing improved. We also investigated PRORP localisation in the mouse inner ear and found high levels in the synapses and nerve fibers that transmit sound. It may be that disruption of RNA processing in the mitochondria of these cells causes hearing loss in this family.

## Introduction

Perrault syndrome (MIM 233400) is a rare autosomal recessive condition characterised by bilateral sensorineural hearing loss (SNHL) affecting both sexes and primary ovarian insufficiency (POI) in 46, XX karyotype females (1). Additional clinical features may be present in some affected individuals, most commonly neurological dysfunction, including: ataxia, hereditary sensory motor neuropathy, nystagmus and mild to moderate intellectual disability (2, 3). Perrault syndrome is challenging to diagnose in males or prepubescent females as both groups are likely to present with non-syndromic SNHL (4). Perrault syndrome is genetically heterogeneous, as causative biallelic variants in six genes have been identified to date: *HSD17B4* (MIM 233400) (5), *HARS2* (MIM 614926) (6), *LARS2* (MIM 615300) (7), *CLPP* (MIM 614129) (8), *C10orf2* (MIM 616138) (9) and *ERAL1* (MIM 607435) (10). These six genes do not account for all cases of Perrault syndrome, suggesting that additional genes remain to be identified (11, 12). Variants in *HSD17B4* are associated with both D-bifunctional protein deficiency, a peroxisomal disorder (13, 14), and Perrault syndrome (5). The other five genes associated with Perrault syndrome are nuclear-encoded mitochondrial proteins and share a common pathology of disrupted mitochondrial protein translation.

We report a novel cause of Perrault syndrome, which shares the common pathology of disrupted mitochondrial translation. Affected individuals in a large consanguineous family are homozygous for a missense variant in *KIAA0391* (MIM 609947), encoding **Pro**teinaceous **R**Nase **P** (PRORP, also known as mitochondrial RNase P protein 3 (MRPP3)), one of the three subunits of the mitochondrial RNase P (mtRNase P) complex. In humans, mtRNase P comprises TRMT10C-SDR5C1-PRORP (also called MRPP1-MRPP2-MRPP3, respectively), and is responsible for cleaving the 5’ end of mitochondrial tRNAs from long polycistronic precursor transcripts (15, 16). We demonstrate that the *KIAA0391* disease-associated variant impairs mtRNase P activity, resulting in the accumulation of unprocessed polycistronic mitochondrial transcripts and leading to a loss of steady-state levels of mitochondrial oxidative phosphorylation (OXPHOS) components. Expression of wild type *KIAA0391* in patient fibroblasts rescued tRNA processing. Finally, immunohistochemistry analyses of mouse auditory sensory epithelium showed prominent levels of PRORP in the efferent synapses and nerve fibres of the auditory hair cells, highlighting a plausible pathology for the sensorineural hearing loss observed in affected individuals.

## Results

### Clinical report

A consanguineous Palestinian family of three affected female siblings, two unaffected female siblings, two unaffected male siblings and their unaffected parents were ascertained (Fig. 1A). All three affected sisters showed absent middle ear acoustic reflex, despite normal tympanometry, when tested in infancy and subsequent audiology examinations in each revealed profound bilateral SNHL (>90 decibels hearing level at all frequencies) (Fig. 1B). The three affected sisters each presented in their late teenage years with primary amenorrhea. Abdominal ultrasound noted the absence of ovaries in all three affected individuals. Hormonal profiles indicated hypergonadotropic hypogonadism (Fig. 1C) with otherwise normal endocrine and biochemical tests and a 46, XX karyotype. The sisters were prescribed estrogen to induce puberty and are currently treated with hormone replacement therapy.

**Figure 1.**
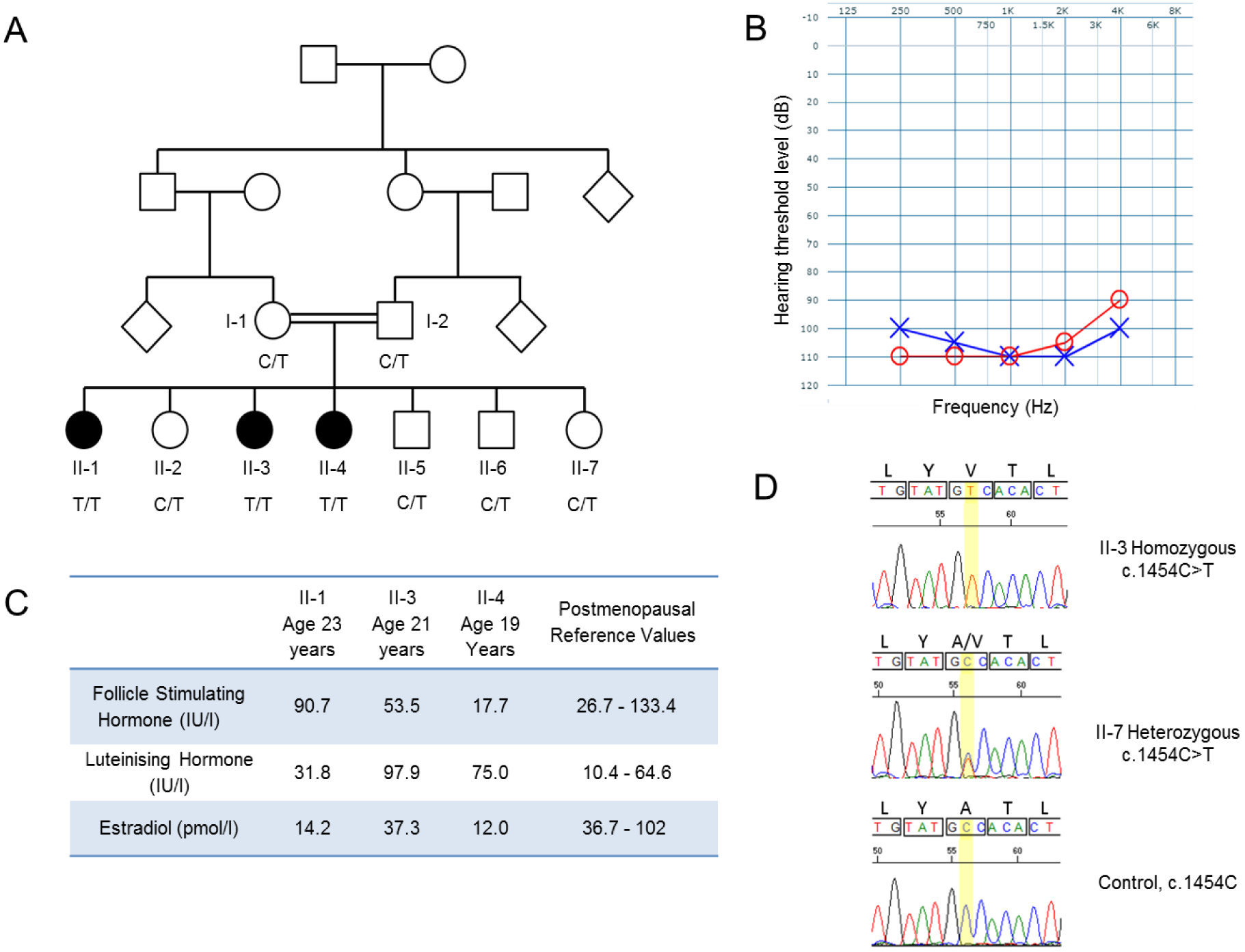
The variant *KIAA0391* c.1454 C>T causes sensorineural hearing loss and primary ovarian insufficiency in a large consanguineous family. **A)** The pedigree for the consanguineous Palestinian family with a variant in *KIAA0391*. The pedigree includes genotypes for the variant *KIAA0391* c.1454 C>T. Filled symbols indicate affected individuals. **B)** Audiogram of affected individual II-4. The hearing level of the left ear is represented by the blue crosses and the right ear by red circles. All three affected sisters show a similar audiometric configuration, with profound hearing loss across all tested frequencies. The hearing threshold level of a normal adult is 0-20 dB (21). **C)** Hormone profiles for the three affected sisters, indicative of hypergonadotropic hypogonadism. Patients with hypergonadotropic hypogonadism have levels of follicle stimulating hormone, luteinising hormone and estrogen in the postmenopausal range (22). **D)** Sanger sequencing traces for an affected and unaffected member of the family and one control sample at position *KIAA0391* c.1454, highlighted in yellow (NM_014672.3).

Each affected sibling has mild intellectual disability, which is not progressive. The sisters were taught in remedial classes but participate in the full activities of daily living, and one sister has paid employment. Echocardiography for each of the affected sisters was normal. All other physical and neurological examinations were normal and the three sisters each have normal height.

### Exome sequencing and variant confirmation

Autozygosity mapping, performed on six siblings, as previously described (17), identified three homozygous regions >2Mb shared between the affected individuals, but not with the unaffected individuals (chromosome 14: 34195478-37228220; chromosome 18: 10090808-12264512; and chromosome 22: 21317876-23416005. Genome build: Hg19.) (18). Whole exome sequencing was performed on one affected individual (II-3). After sequence variants in the autozygous regions were filtered to remove variants seen more than once in >800 previously sequenced exomes, three variants remained. Two variants with a minor allele frequency of greater than 1% in the Exome Variant Server (19) and dbSNP frequency (20) respectively were excluded, as they were too common to result in a rare inherited disorder. One variant was novel, *KIAA0391* c.1454C>T; p.(Ala485Val) (Genbank: NM_014672.3).

*KIAA0391* encodes the endonuclease subunit of the mtRNase P complex, PRORP, which catalyses Mg^2+^-dependent phosphodiester-bond cleavage of 5’ extensions of mitochondrial tRNAs (15). Residue Ala485 is highly conserved in PRORP from multiple species (Supplemental Fig. S1). The p.Ala485Val variant is predicted to be deleterious by *in silico* pathogenicity tools, including SIFT (23), PolyPhen2 (24) and MutationTaster (25), and was absent in 100 ethnically matched controls, in the ExAC server (26), dbSNP (20) or EVS (19) – comprising a minimum of 70,000 individuals. Of note, homozygous loss of function variants in *KIAA0391* are absent from publically available databases noted above and a cohort of >3,200 British Pakistani individuals (27). Segregation of the homozygous variant with the Perrault syndrome phenotype was confirmed via Sanger sequencing (Fig. 1A and 1D). Sequence analysis of *KIAA0391* in five unrelated individuals with Perrault syndrome (11), without a disease-causing variant in the known Perrault syndrome genes, did not identify putative pathogenic variants in a second family.

### Patient cells contain normal levels of mtRNase P but display decreased levels of oxidative phosphorylation (OXPHOS) components containing mitochondria-encoded proteins

We investigated the steady-state levels of TRMT10C, SDR5C1 and PRORP in patient fibroblasts by Western blot analysis and found no decrease in the levels of the mtRNase P subunits compared to controls (Fig. 2A), suggesting that the p.(Ala485Val) variant does not markedly influence the stability of PRORP or the other mtRNase P subunits. We detected decreased steady-state levels of subunits of respiratory chain complex I (NDUFB8) and complex IV (COXI and COXII) subunits in patient fibroblasts compared to controls – both complexes containing mitochondrial DNA-encoded subunits - with no change of other OXPHOS components observed, most notably complex II, which is entirely nuclear-encoded(Fig. 2B). These results indicate a generalised defect of mitochondrial translation in the patient fibroblasts.

**Figure 2.**
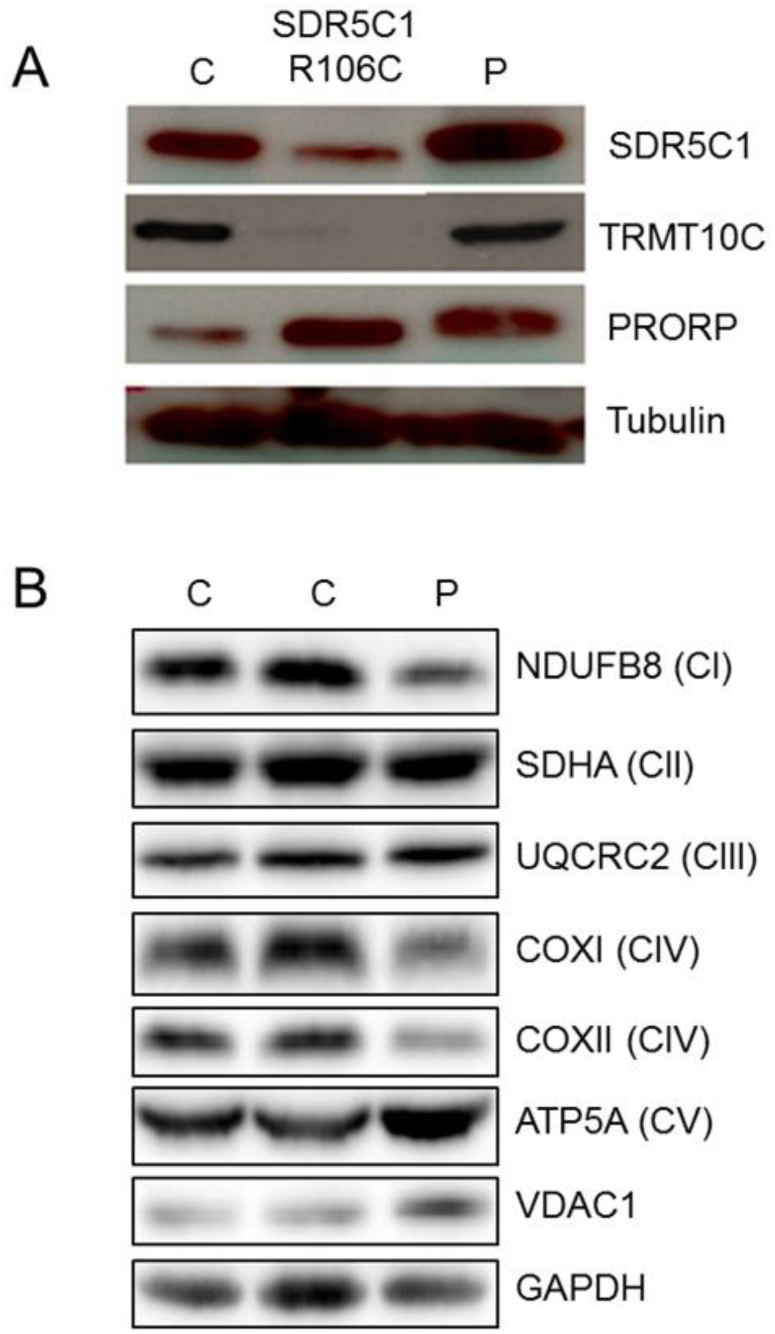
Patient fibroblasts show no reduction in subunits of mtRNase P, but show reduced levels of mitochondrial DNA encoded OXPHOS subunits. **A)** Western blot analysis of mtRNAse P subunits TRMT10C, SDR5C1 and PRORP in fibroblasts from a healthy control (C); an individual with the R106C pathogenic substitution in SDR5C1 (28) and individual II-4 with the p.Ala485Val variant in PRORP (P). Steady state levels of both TRMT10C and SDR5C1 are known to be reduced in patients with HDS10 disease due to the known pathogenic variant in SDR5C1, R106C (28). **B)** Western blot analysis of proteins of the five oxidative phosphorylation complexes. Included are two control samples (C) and a patient sample (P), II-4. VDAC1 serves as a mitochondrial loading control.

### Patient cells display disrupted mitochondrial RNA processing

The levels of unprocessed mitochondrial transcripts in fibroblasts from individual II-4 were assessed by Northern blot analysis. An *MT*-*ND1* probe detected an RNA transcript of approximately 2.5 kb in the patient sample not observed in the controls suggesting impaired 5’-end processing of tRNA^Leu(UUR)^ (Fig. 3). A larger RNA species appeared on a longer exposure for the *MT*-*ND1* probe indicating that tRNA^Val^ processing is also decreased. An *MT*-*ND2* probe detected a large RNA transcript in the patient sample at approximately 4 kb indicating impaired 5’-end processing of tRNA^Ile^, tRNA^Met^ and tRNA^Trp^. A longer exposure revealed multiple bands, indicating the accumulation of transcripts due to the impaired 5’-end processing at multiple tRNA sites. An *MT*-*CO2* probe showed a transcript of approximately 1.7 kb not detected in the controls suggesting impaired processing of tRNA^Lys^. Multiple bands in a longer exposure using the *MT*-*CO2* probe indicated decreased processing across multiple tRNA sites. An *MT*-*ND6* probe detected a transcript at approximately 2.3 kb in the patient sample, which can be explained by impaired 5’ processing of tRNA^Glu^. Taken together, the accumulation of various large unprocessed mitochondrial RNA transcripts across multiple tRNA sites indicates a generalised insufficiency of mitochondrial tRNA processing.

**Figure 3.**
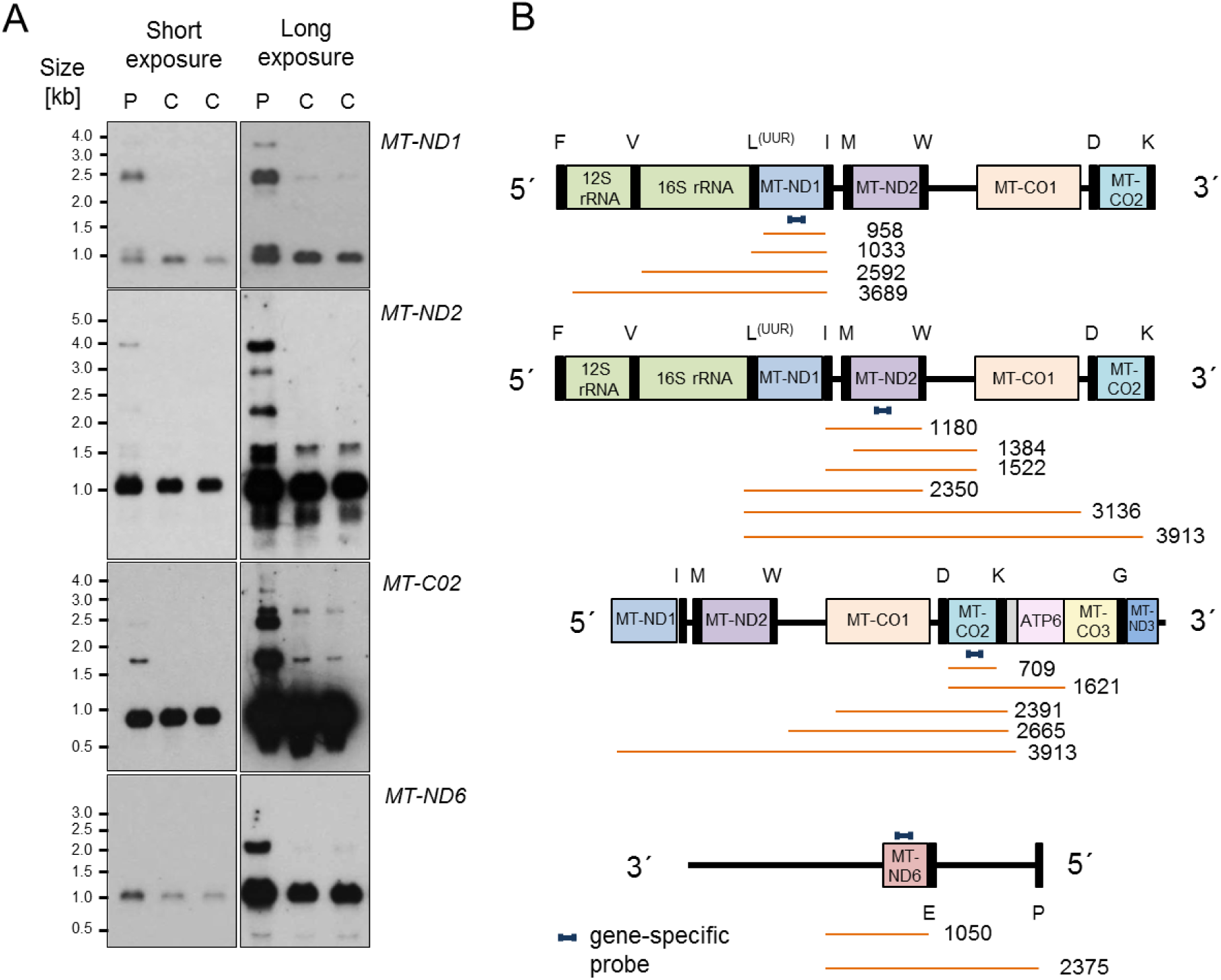
Impaired mitochondrial RNA processing is detected in patient fibroblasts. **A)** Northern blot assessment of RNA extracted from patient (P) fibroblasts (II-4) and two control (C) samples using strand specific probes designed to complement four different mitochondrial gene transcripts; *MT*-*ND1*, *MT*-*ND2*, *MT*-*CO2* and *MT*-*ND6.* A long and short exposure of the blots is shown. **B)** Schematic representations of mitochondrial genome regions, the probes (dark blue) and expected fragment sizes in bp (orange) are shown to the right of each blot.

### The PRORP p.Ala485Val disease variant displays decreased mtRNase P activity

The disease-associated variant p.Ala485Val is situated in the metallonuclease domain of PRORP, close to four conserved aspartate residues implicated in metal ion binding (Fig. 4A and 4B) (29). Replacing the conserved alanine at residue 485 with the bulkier valine could distort the active site and impair catalysis by interfering with proper coordination of the metal ions, thereby reducing the endonucleolytic activity of PRORP. To determine whether the p.Ala485Val variant alters the catalytic activity of PRORP, we compared the enzymatic activity of the disease-associated isoform and the wild type protein in the RNase P complex.

**Figure 4.**
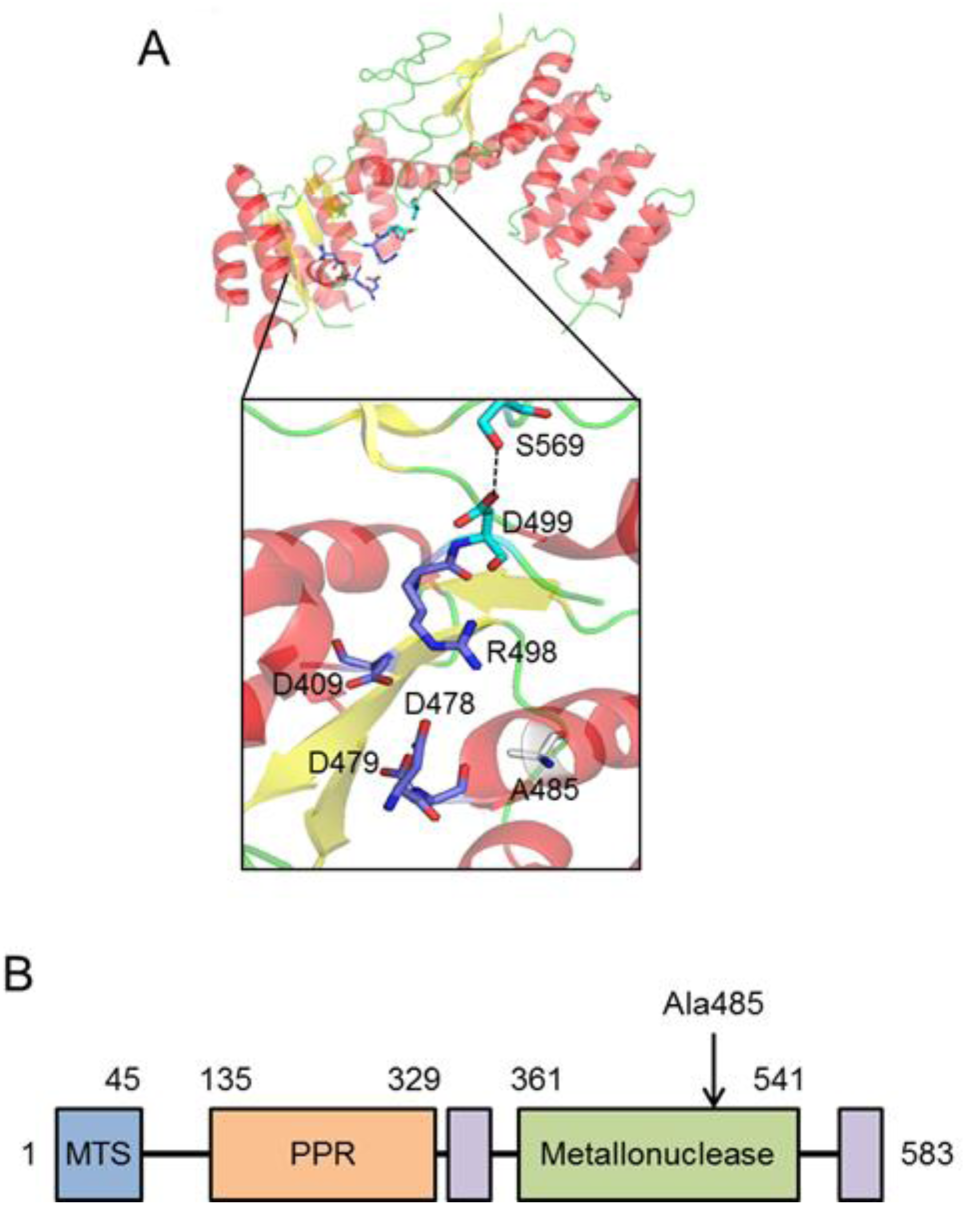
The residue Ala485 is situated in the metallonuclease domain of PRORP close to conserved residues. **A)** The protein structure of human PRORP; the enlarged region is part of the metallonuclease domain. Residue Ala485 is situated in the centre of the active site close to the conserved aspartate residues that coordinate catalytic magnesium ions. **B)** A bar diagram of the domains of human PRORP (30). The location of the variant residue Ala485 is noted on the diagram. Mitochondrial targeting sequence, MTS; pentatricopeptide repeat domain, PPR.

The three wild type proteins of the mtRNase P complex TRMT10C, SDR5C1 and PRORP as well as the PRORP p.Ala485Val variant protein were individually produced by recombinant expression in *E.coli* and purified. During the assessment of the recombinant protein PRORP p.Ala485Val we noted that it had a higher melting temperature than wild type PRORP (Supplemental Fig. S2) suggesting reduced flexibility of the variant PRORP. Recombinant mtRNase P was reconstituted *in vitro* and 5’ end processing assays were performed with radiolabelled precursor tRNA (pre-tRNA) (Fig. 5 and Supplemental Fig. S3). Assays were performed with three mitochondrial tRNA substrates. Variants in *HSD17B10* (encoding SDR5C1) causing HSD10 disease (MIM 300438) were reported to affect the accumulation of unprocessed transcripts from the heavy strand, but not those from the light strand (28). We therefore selected the heavy strand-encoded pre-tRNA^Ile^ and the light strand-encoded tRNA^Tyr^. Additionally, Pre-tRNA^His-Ser(AGY)-Leu(CUN)^ was selected, because variants in *HARS2* and *LARS2*, which encode mitochondrial histidyl and leucyl aminoacyl synthetases, respectively, cause Perrault syndrome (6, 7).

**Figure 5.**
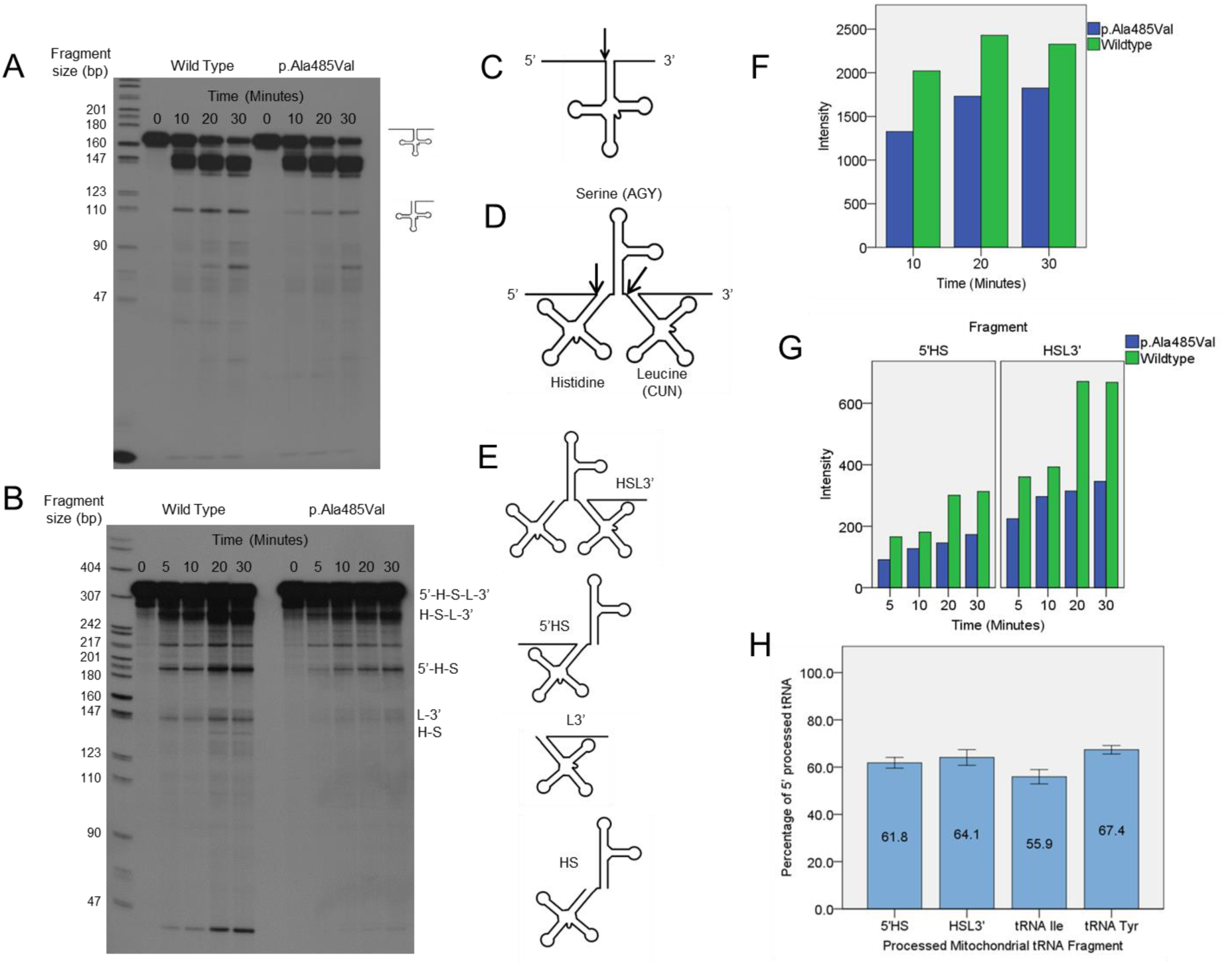
*In vitro* mtRNase processing assays show the variant p.Ala485Val PRORP produces significantly less 5’ end processed tRNA than wild type PRORP. **A)** Mitochondrial pre-tRNA^Tyr^ was subjected to cleavage by reconstituted recombinant mtRNase P containing either wild type or p.Ala485Val PRORP. Aliquots were removed from the reactions and stopped at the time points indicated, and resolved by urea-PAGE. **B)** Mitochondrial pre-tRNA^His-ser(AGY)-Leu(cuN)^ was subjected to cleavage by reconstituted recombinant mtRNase P containing either wild type or p.Ala485Val PRORP. Processing as in **(A)** with an additional time point added at 5 minutes. See **(E)** for fragment labelling. **C)** A simplified drawing of pre-tRNA; the arrow indicates the site of mtRNase P processing. **D)** A drawing of pre-tRNA^His-Ser(AGY)-Leu(CUN)^, (31). The arrows indicate the sites of mtRNase P processing. **E)** The fragments produced from pre-tRNA^His-Ser(AGY)Leu-(CUN)^ by mtRNase P processing listed in order of descending size; the 5’-end fragment not included. **F)** Quantitative analysis of pre-tRNA^Tyr^ processing experiment shown in **(A)**. There was significant difference between the p.Ala485Val and wild type mtRNaseP over three replicate experiments (n = 9 paired wild type and equivalent variant time points across 3 replicate experiments, P<0.01). **G)** Quantitative analysis of pre-tRNA^His-Ser(AGY)-Leu(CUN)^ processing experiment shown in **(B)**, only the fragments HSL3’and 5’HS are shown. The HSL3’ fragment at 30 minutes shows reduction compared to the fragment at 20 minutes likely because of further processing to produce the HS fragment. There was significant difference between the p.Ala485Val and the wild type mt RNaseP (HSL3’; n = 12 paired time points across 3 replicate experiments, P< 0.01 and 5’HS; n = 12 paired time points across 3 replicate experiments, P< 0.01). **H)** The average tRNA processing of the four major fragments from the three substrates as compared to the wild type processing. Wild type is set to 100%. Data presented as the mean of 3 replicate experiments (all time points included) +/− SEM.

Since mtRNase P is an endonuclease, two fragments result from the cleavage of pre-tRNA^Tyr^ and pre-tRNA^Ile^: the removed 5’-leader and the 5’-mature tRNA (Fig. 5A and 5C; Supplemental Fig. S3A). From the pre-tRNA transcript containing three tRNAs, the combination of two mtRNase P sites produces five possible fragments (31) (Fig. 5B, 5D and 5E). The variant mtRNase P complex generated markedly less processed tRNA than the wild type complex across all time points and reactions. Quantitative phosphorimaging revealed a diminution of cleavage products by ~35-45% (p<0.01) depending on the pre-tRNA substrate used (Fig. 5F, 5G, 5H, and Supplemental Fig. S3B). There was no significant difference in the reduction of 5’-end processing by PRORP p.Ala485Val-containing mtRNase P between the different pre-tRNA substrates analysed (p>0.05), consistent with the accumulation of both heavy and light-strand transcripts seen in patient fibroblasts (Fig. 3).

### tRNA processing in patient cells can be rescued by expression of wild type *KIAA0391*

Rescue experiments were used to connect the accumulation of unprocessed RNA transcripts seen in the patient cells to the variant PRORP p.(Ala485Val). Patient fibroblasts were transfected with a plasmid containing the wild type *KIAA0391* sequence. Unprocessed RNA transcripts were quantified by qPCR across processing sites (Fig. 6A). In untransfected patient cells there was a greater than 6-fold increase in both pre-tRNA^Leu^ and pre-tRNA^Val^ transcripts in comparison to controls. When the patient cells were transfected with wild type PRORP, this increase was reduced to about 3-fold for pre-tRNA^Leu^, and pre-tRNA^Val^ levels came back close to the control. Transfection efficiency, which is 70% maximum, could account for tRNA processing not achieving wild type levels in transfected patient cells.

**Figure 6.**
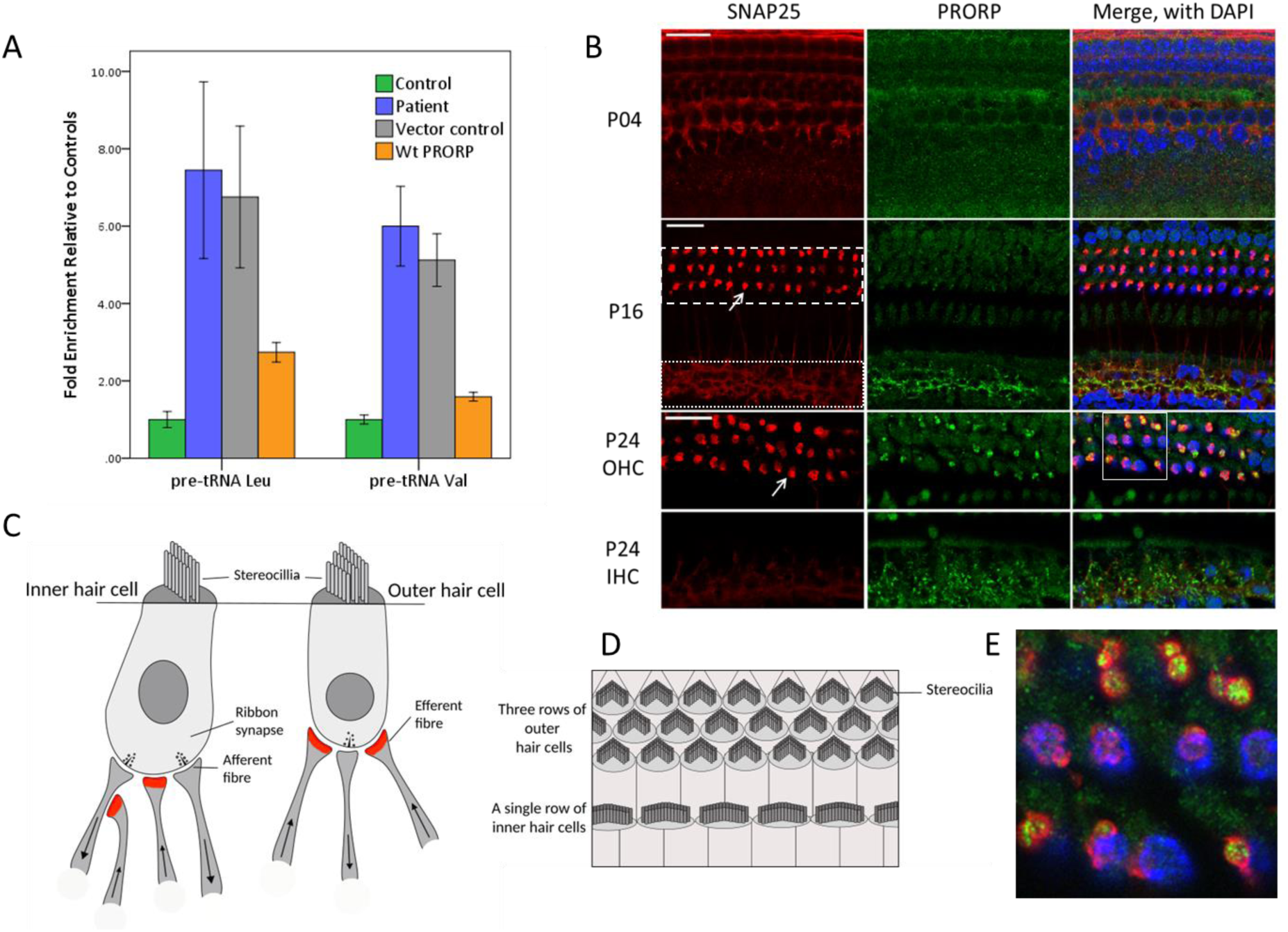
Expression of wild type *KIAA0391* encoding PRORP rescues tRNA processing and localisation in the mouse organ of Corti shows high levels of PRORP in the synapses and nerve fibres of hair cells. **A)** Mitochondrial precursor transcripts containing tRNA^Val^ and tRNA^Leu^ were quantified in control and patient fibroblasts using real time PCR. The patient samples are presented as a relative fold increase of the relevant controls. The precursor transcripts in patient samples were measured in non-transfected cells, cells transfected with the empty vector and in cells transfected with the vector containing the wild type *KIAA0391* sequence. For the control samples non-transfected fibroblasts were quantified. Fibroblast cell cultures of three different healthy donors served as controls. (mean ± SEM). Untransfected patient cells showed a greater than 5 fold increase in both pre-tRNA^Leu^ and pre-tRNA^Val^ transcripts compared to controls. Transfection of patient cells with wild type *KIAA0391* reduced precursor transcripts by ~75% for pre-tRNA^Leu^ and ~ 65% pre-tRNA^Val^, rescuing tRNA processing. **B)** Confocal fluorescence microscopy optical sections of the whole mount organ of Corti samples from C57/BJ6 mice at postnatal days 4, 16 and 24 (P4,P16 and P24, correspondingly) showing localization of PRORP protein (green). Samples were counterstained with DAPI (nuclear DNA marker, blue) to visualize the nuclei of hair cells and SNAP25 (presynaptic membrane marker, red) to visualize efferent synapses at the base of OHCs and nerve fibres and synaptic buttons at the base and around IHCs. In the panels for P4 and P16 the dashed white line outlines the area of the outer hair cells (OHC) at the nuclear level and the dotted white line outlines the area around the inner hair cell nuclei (IHC). The white arrows point to one of the OHC efferent presynaptic buttons. The two panels at P24 represent two optical sections through the same organ of Corti sample at different focal plains to visualize OHC synaptic area (top) and IHC synaptic area (bottom). The scale bar is 20μm. **C)** An illustration of the innervation of inner hair cells and outer hair cells. Arrows in the nerve fibres indicate the direction of transmission. The area of SNAP25 staining is shown in red. **D)** An illustration of the arrangement of hair cells in the organ of Corti in the same orientation as shown in Panel **B. E)** An enlarged view of the area inside the white box in Panel **B** showing the co-localisation of SNAP25 (red) and PRORP (green) with the OHC nuclei shown in blue.

### PRORP is present in hair cell synapses and neurons of the mouse organ of Corti

To understand why the variant in PRORP, p.Ala485Val, could be associated with hearing loss in the affected family, we undertook localisation studies of PRORP in the mouse organ of Corti. The organs of Corti from C57/BJ6 mice at postnatal (P) day 4 (P4), P16 and P24, were immunofluorescently stained to reveal the localisation of endogenous PRORP. The samples were also counterstained with a presynaptic membrane marker SNAP25 (synaptosome-associated protein 25), and DAPI, a fluorescent stain for double-stranded DNA that highlights the cell nucleus.

In mice, the onset of hearing occurs at approximately P12-P14 (32). At P4, the organ of Corti is not fully mature and low levels of SNAP25 protein are detected at presynaptic hair cell membranes (Figure 6b) (33). At P4, diffuse PRORP staining at low levels is observed. Shortly after the onset of hearing at P16, the outer hair cell (OHC) efferent synapses and the inner hair cell (IHC) nerve fibres are highlighted by SNAP25. At P16 PRORP signal is detected around the base of inner hair cells where afferent and efferent synapses are situated. At the same time, we observed low levels of PRORP in the hair cell bodies and in the efferent synapses of OHCs. By P24, PRORP immunoreactivity is increased at the efferent synapses of OHCs and around the base of inner hair cells. PRORP partially co-localises with SNAP25 staining around IHCs (see P16 images), suggesting that PRORP could be present not only in the mitochondria of efferent synapses but possibly afferent synapses and nerve fibres around the IHCs. Low levels of PRORP signal is present in OHC bodies at P24 alongside the more intense signal seen at the OHC synapses and IHC base. At P24, the mitochondrial marker Tom20 labels mitochondria mainly within hair cell bodies and does not show increased signal in mitochondria within the supporting cells or efferent synaptic buttons (Supplementary Fig. S4), arguing for structurally different mitochondria with different protein compositions present in different cell types (34, 35). It is likely that the high levels of PRORP found in the subset of mitochondria associated with the synapses and neurons of the organ of Corti hair cells reflect those cells’ increased demand for genes involved in mitochondrial tRNA processing and translation.

## Discussion

In this report we provide evidence that Perrault syndrome is caused by a novel homozygous variant in *KIAA0391*, a gene not previously implicated in this disorder. *KIAA0391* is the nuclear gene encoding PRORP, the catalytic subunit of the mtRNase P complex (15). *In vitro* tRNA processing assays supported the *in silico* predictions that the amino acid substitution in PRORP p.Ala485Val, the expected result of the nucleotide alteration c.1454C>T, would be damaging by impairing the processing of mitochondrial tRNAs. The absence of variants in *KIAA0391* in other known cases of Perrault syndrome suggests that variants in *KIAA0391* are not a common cause of this heterogeneous condition. However, our discovery provides further important insights into the pathogenesis of hearing loss and ovarian insufficiency. The functional work reported here indicates the mitochondrial respiratory chain deficiency seen in this family can be attributed to a variant in *KIAA0391*. An independent family would provide additional data linking variants in *KIAA0391* with Perrault syndrome as would recapitulation of the Perrault syndrome phenotype in an animal model of the p.Ala485Val substitution in PRORP.

The mtRNase P complex is composed of three proteins, TRMT10C-SDR5C1-PRORP, each encoded by the nuclear genome and post-translationally imported into the mitochondria matrix via an N-terminal mitochondrial targeting sequence (15, 16). Mitochondrial tRNAs are processed at the 5’ end by mtRNase P (15) and at the 3’ end by mtRNase Z, encoded by *ELAC2* (36, 37). This tRNA processing excises most of the RNA species from the polycistronic mitochondrial precursor transcripts according to the tRNA punctuation model (38, 39).

We observed normal levels of PRORP in patient fibroblasts (Fig. 2A), which contrasts with the reduction of the other two subunits of mtRNase P, SDR5C1 (28, 40) and TRMT10C (41) in patients with inherited deficiencies of those proteins. Accumulation of multiple unprocessed transcripts in patient fibroblasts indicated a generalised defect in mitochondrial tRNA processing (Fig. 3), leading to a mild but observable downstream effect on mitochondrial protein synthesis as evidenced by decreased steady-state levels of complex I and complex IV subunits (Fig. 2B). A similar, but more pronounced, OXPHOS defect was also observed in patients with pathogenic variants in *TRMT10C* (41) and *HSD17B10* (encoding SDR5C1) (42). OXPHOS deficiency has also been associated with Perrault syndrome in patients with variants in *ERAL1* (10). The pattern of OXPHOS deficiency in patients with Perrault syndrome due to *ERAL1* variants is similar to that seen in our patient, suggesting that they may share a common pathogenesis.

Biallelic variants, which are predicted to result in complete loss of function, have not been identified in any of the three subunits of the mtRNase P complex. All three mtRNase P complex genes are among the core set of genes essential for the survival of cells in tissue culture (43, 44). There are no knockout mouse models for *trmt10c* and the homozygous knockout of *hsd17b10* results in an early embryonic lethal phenotype (45). Knockout of *1110008L16Rik* (*KIAA0391* homolog) is embryonic lethal, but cardiac and skeletal muscle specific *1110008L16Rik* conditional knockout mice have been produced (46). These mice die at 11 weeks due to cardiomyopathy. The conditional knockout mice showed a significant reduction in the synthesis of mitochondrial encoded proteins with no alteration in nuclear encoded proteins. The reduction in mitochondrial-encoded proteins was accompanied by a significant reduction in mitochondrial respiration (46). The mice also showed an accumulation of unprocessed mitochondrial RNA transcripts, but in contrast to patients with defects in subunits of mtRNase P, the mice had no mature transcripts (46). The lack of mature mitochondrial transcripts may be the result of the complete loss of PRORP protein function in mice in comparison to patients with predicted hypomorphic variants in *TRMT10C, HSD17B10* or *KIAA0391*.

Despite the similarities in defective mitochondrial tRNA processing, variants in the three subunits of mtRNase P result in markedly differing clinical phenotypes. Pathogenic variants in *TRMT10C* cause a lethal, childhood multisystem disorder characterised by muscular hypotonia, SNHL and metabolic acidosis (41). Pathogenic variants in *HSD17B10*, encoding SDR5C1 (protein also known as HSD10, HADH2, MRPP2 or ABAD), cause HSD10 disease, which manifests as a severe, infantile-onset neurodegenerative condition with cardiomyopathy (40, 47). The clinical presentation of Perrault syndrome is less severe than of individuals with TRMT10C (41) and SDR5C1-associated diseases (40). The tissue specific phenotype of Perrault syndrome (SNHL and POI) in comparison to the more systemic presentations of TRMT10C- and SDR5C1-associated phenotypes may be accounted for by the differing function of the proteins. TRMT10C and SDR5C1 are also important for methylation at position 9 of mitochondrial tRNAs, which stabilises the tertiary L-shaped structure of some tRNAs (16),whereas PRORP is required for the nuclease function of mtRNase P only (15, 16). It is possible that more deleterious variants in *KIAA0391* would result in further reduction in PRORP activity and a more severe multisystem disorder, similar to that seen in individuals with *LARS2* variants (48). Conversely, hypomorphic alleles in *KIAA0391* could result in milder degrees of SNHL or POI only.

Immunohistochemistry of the mouse organ of Corti revealed the localisation of PRORP in hair cells with higher levels associated with the efferent synapses of the outer hair cells and nerve fibres of the inner hair cells. This particular pattern of PRORP localisation becomes apparent after the onset of hearing and involves only mitochondria of some cell types. It seems likely that a subpopulation of mitochondria associated with neurons in the organ of Corti are enriched for PRORP and may require a higher level of mitochondrial translation, after hearing onset.

Recently, the expression levels of hearing loss genes were examined in different areas of the mouse cochlea. Four of the six genes associated with Perrault syndrome were examined, and it was found that they showed higher expression levels in the spiral ganglion neurons than in other parts of the cochlea, including the organ of Corti (49). This pattern of expression correlates with our immunohistochemistry results, as the cell bodies of the nerve fibres with efferent synapses that terminate at the base of OHC and IHC are in the brainstem, and IHC afferent synapses and their nerve fibres originate from the spiral ganglion neurons. Dysfunction of PRORP causing disruption of the hair cell efferent or afferent signalling, in conjunction with malfunctioning mitochondria of hair cell bodies, could be the pathological mechanism behind the hearing loss in the family reported here. The gene expression patterns from other Perrault syndrome genes suggest that they may share a similar pathology of neuronal origin with the case presented here (49). Our results together with the results from Nishio *et al* (49) suggest that the neurons in the spiral ganglion and brainstem may have an increased level of mitochondrial translation, perhaps making them vulnerable to disruption of this pathway during sustained sound stimulations after the onset of hearing.

Pathogenic variants resulting in distinct clinical phenotypes have now been identified in all the genes encoding subunits of mtRNase P. It is currently unclear why pathogenic variants in the different subunits of mtRNase P cause distinct clinical phenotypes. Investigation into this could provide insights into differences in the pathogenic mechanisms.

Most genes previously associated with Perrault syndrome (*HARS2*, *LARS2*, *C10orf2*, *CLPP* and *ERAL1*) encode proteins implicated in mitochondrial protein translation (6-10). Our finding that biallelic variants in *KIAA0391*, encoding PRORP, result in impaired mitochondrial tRNA processing expands the spectrum of genes that can cause Perrault syndrome and provides additional evidence that Perrault syndrome is caused by impaired mitochondrial translation. Further, this discovery lends additional support to the hypothesis that genes involved in mitochondrial translation are candidates for genetically unresolved cases of Perrault syndrome.

## Materials and Methods

### Ethical approval

All patients provided written informed consent in accordance with local regulations. Ethical approval for this study was granted by the National Health Service Ethics Committee (16/WA/0017) and University of Manchester. The NIH Animal Use Committee approved protocol 1263-15 to T.B.F. for mice.

### Autozygosity mapping and whole exome sequencing

Autozygosity mapping was performed on six members of the family (II-1, II-2, II-3, II-4, II-6 and II-7) using the Affymetrix Genome-wide SNP6.0 arrays as previously described (17). Whole exome sequencing was performed on DNA extracted from lymphocytes from individual II-3. The Agilent SureSelect Human All Exon V5 Panel was used for library preparation and sequencing was performed on the HiSeq 2500 (Illumina) as previously described (50). The variant *KIAA0391* c.1454C>T; p.(Ala485Val) has been submitted to the relevant LOVD database (http://www.lovd.nl/3.0/home).

### Confirmation of variants

Variants were confirmed in the family via Sanger sequencing using the ABI big Dye v3.1 (ThermoFisher) sequencing technology. Primer sequences are available on request.

### Assessment of protein and RNA levels in patient fibroblasts

Western blot analyses for the mtRNase P subunits were performed as previously described (28) using cell lysates from dermal fibroblasts (II-4). 30 mg of protein was separated on a 10% SDS-polyacrylamide gel and transferred to a polyvinylidene difluoride membrane (GE Healthcare). Membranes were blocked and subsequently incubated with the primary antibodies (28) and detected with HRP-conjugated secondary antibodies (Dako). The blots were developed using the ECL Western Blotting Analysis Detection system (GE Healthcare).

Western blots for the respiratory chain complexes (n=3) were performed using fibroblast cell lysates (II-4). Cell lysates were incubated with sample dissociation buffer, separated by 12% SDS–PAGE and immobilized by wet transfer on to PVDF membrane (Immobilon-P, Millipore Corporation). Proteins of interest were bound by overnight incubation at 4°C with primary antibodies followed by HRP-conjugated secondary antibodies (Dako Cytomation) and visualized using ECL-prime (GE Healthcare) and BioRad ChemiDoc MP with Image Lab software. Antibody details are available in the Supplemental Materials.

Northern blot analysis was performed as previously described (28). The NorthernMax kit from Ambion was used. 2–5 μg of total RNA, from fibroblasts (II-4), was separated on a 1% denaturing agarose gel. RNA was then transferred to nylon membrane (Hybond-N + Amersham, GE Healthcare) by capillary transfer, UV cross-linked and subjected to hybridization with biotinylated probes. Signals were detected using the BrightStar BioDetect kit (Ambion). A biotinylated RNA size marker (BrightStar RNA Millenium Marker, Ambion) was used to determine the size of RNA species. Probe sequences as previously described (28).

### Preparation of the PRORP p.Ala485Val sequence for bacterial expression

The plasmid pET28-b(+) containing the coding sequence for PRORP (15) was mutagenized as previously described (51) with the synthetic oligonucleotide PRORP_p.Ala485Val (see Supplemental data for oligonucleotide sequence). The potential mutagenized plasmids were extracted using the GenElute HP Plasmid miniprep Kit (Sigma Aldrich) and the variant was confirmed by DNA sequencing.

### Recombinant expression and purification of TRMT10C, SDR5C1 and PRORP (wt and p.Ala485Val)

The templates for recombinant expression of *TRMT10C*, *SDR5C1 and PRORP* were as detailed in (15) and above. Expression was induced in *E.coli* Rosetta2 DE3 (Novagen) using Overnight Express medium (Novagen). Affinity chromatography of the His-tagged proteins was performed as previously described (15). Their purity was assessed by SDS-PAGE. Aliquots of purified proteins were dialysed overnight at 4°C in 20 mM Tris-Cl pH 7.4, 100 mM NaCl, 15% glycerol, then flash frozen and stored at −80°C.

### Preparation of mitochondrial pre-tRNA transcripts

The templates for pre-tRNA^Tyr^ and pre-tRNA^Ile^ were as described in (15). The template for the pre-tRNA^His-Ser(AGY)-Leu(CUN)^ was as described in the Supplemental data. Run off *in vitro* transcription to produce body labelled pre-tRNA substrate was performed as previously described (52).

### Pre-tRNA processing assays

Pre-tRNA processing assays were performed as previously described (15, 53). The TRMT10C, SDR5C1 and PRORP proteins were mixed in a 2:4:1 molar ratio as described (15). 6% (w/v) acrylamide 8M urea gels were used to resolve substrate and cleavage products. Dried gels were exposed to X-ray film at −80°C or exposed to a phosphorimaging screen to detect assay precursors, intermediates and products. See Supplemental Materials for further details.

### Real-time PCR

Real-time PCR and analysis was performed as previously described (28). Total RNA was isolated from patient and control dermal fibroblasts and treated with DNase. Reverse transcription was performed using tRNA specific primers containing an adaptor sequence for subsequent real-time PCR. Primers sequences are as previously described (28). Real time PCR was performed using standard conditions (60°C elongation, 40 cycles) and relative expression of precursor transcripts in patient fibroblasts was calculated using the ct value for the appropriate transcript from three different healthy controls. Ubiquitin B expression served as the reference for both patient samples and controls.

### Rescue experiments

The coding sequence of wild type *KIAA0391* was cloned in an expression vector as described previously (28). The coding sequence was verified by Sanger sequencing. Patient dermal fibroblasts (2.0 × 10^5^) were seeded in 25 cm^2^ flasks and cultivated in 4 ml MEM medium. The next day cells were transfected using 3μg plasmid DNA (PRORP-vector and empty vector as transfection control) and 7.5 μl Turbofect transfection reagent and cultured for additional 48 hours. The cells were then harvested and precursor tRNAs were quantified by real-time PCR as described previously (28).

### Localisation of PRORP in the mouse organ of Corti

The NIH Animal Use Committee approved protocol 1263-15 to T.B.F. for mice. C57/BJ6 mice at ages P4, P16 and P24 were euthanised, the cochlear capsule was removed and fixed with 4% paraformaldehyde in PBS for 2 hours. The samples were microdissected and the organ of Corti was permeabilised with 1% Triton X-100 in PBS for 30 min followed by three 10 min washes with 1X PBS. Nonspecific binding sites were blocked with 5% normal goat serum and 2% BSA in PBS for 1 h at room temperature. Samples were incubated for 2 h with rabbit polyclonal PRORP antibody (MRPP3, Proteintech - Catalog number: 20959-1-AP) at 1μg/ml and mouse monoclonal SNAP25 antibody (Santa Cruz, sc-136267) at 1μg/ml followed by several rinses with PBS. Samples were incubated with goat anti-rabbit IgG Alexa Fluor 488 conjugated secondary antibody and goat anti-mouse Alexa Fluor 568 conjugated secondary antibody (Molecular Probes) for 30 min. Samples were washed several times with PBS and were mounted with ProLongGold Antifade staining reagent with DAPI (Molecular Probes) and examined using LSM780 confocal microscope (Zeiss Inc) equipped with 63X, 1.4 N.A. objective.

### Statistical analysis

The Wilcoxon paired test (N=<20) was used with a significance value of 0.05 to test for differences between the wild type and p.Ala485Val pairs for each pre-tRNA substrate.

To test for differences in the percentage reduction in the tRNA output of the p.Ala485Val variant between substrates the data was determined to be normally distributed using the Shapiro-Wilk test. One-way ANOVA with a significance value of 0.05 was used to test for differences.

## Acknowledgements

We would like to thank the family for their participation.

## Author Contributions

Conceptualisation, I.H., W.G.N. and R.T.O.; Methodology, W.G.N., R.T.O., L.A.M.D., I.H., J.O., A.J.D., W.R., A.A., S.O., W.W.Y., J.Z., I.B., T.B.F., and R.W.T.; Software, S.G.W. and S.S.B.; Formal Analysis, S.G.W., S.S.B., L.A.M.D., J.E.U. and J.O.; Investigation, W.G.N, R.T.O., L.A.M.D., I.H., J.E.U, J.O., A.J.D., A.A., S.D, N.A., Z.B., S.Y., S.O., K.T, W.W.Y., J.Z., I.A.B, M.B., and R.W.T.; Writing - original draft, L.A.M.D and W.G.N; Writing - review and additional input, all; Visualisation, I.H., L.A.M.D, A.J.D., A.A., Z.B., S.O., K.T, W.W.Y., J.Z., R.W.T., I.B. and R.T.O.; Supervision, K.J.M, T.B.F, W.G.N. and R.T.O.

## References

1. Pallister PD, Opitz JM. The Perrault syndrome: autosomal recessive ovarian dysgenesis with facultative, non-sex-limited sensorineural deafness. Am J Med Genet. 1979;4(3):239–46.

2. Gottschalk ME, Coker SB, Fox LA. Neurologic anomalies of Perrault syndrome. Am J Med Genet. 1996;65(4):274–6. Epub 1996/11/11.

3. Jenkinson EM, Clayton-Smith J, Mehta S, Bennett C, Reardon W, Green A, et al. Perrault syndrome: Further evidence for genetic heterogeneity. J Neurol. 2012;259(5):974–6.

4. Newman WG, Friedman TB, Conway GS. Perrault Syndrome. University of Washington, Seattle; 2014 [updated 2014/09/25; cited 2014 17/11]; Available from: http://www.ncbi.nlm.nih.gov/books/NBK242617/.

5. Pierce SB, Walsh T, Chisholm KM, Lee MK, Thornton AM, Fiumara A, et al. Mutations in the DBP-Deficiency Protein HSD17B4 Cause Ovarian Dysgenesis, Hearing Loss, and Ataxia of Perrault Syndrome. Am J Hum Genet. 2010;87(2):282–8.

6. Pierce SB, Chisholm KM, Lynch ED, Lee MK, Walsh T, Opitz JM, et al. Mutations in mitochondrial histidyl tRNA synthetase HARS2 cause ovarian dysgenesis and sensorineural hearing loss of Perrault syndrome. Proc Natl Acad Sci U S A. 2011;108(16):6543–8.

7. Pierce SB, Gersak K, Michaelson-Cohen R, Walsh T, Lee MK, Malach D, et al. Mutations in LARS2, encoding mitochondrial leucyl-tRNA synthetase, lead to premature ovarian failure and hearing loss in Perrault syndrome. Am J Hum Genet. 2013;92(4):614–20.

8. Jenkinson EM, Rehman AU, Walsh T, Clayton-Smith J, Lee K, Morell RJ, et al. Perrault syndrome is caused by recessive mutations in CLPP, encoding a mitochondrial ATP-dependent chambered protease. Am J Hum Genet. 2013;92(4):605–13.

9. Morino H, Pierce SB, Matsuda Y, Walsh T, Ohsawa R, Newby M, et al. Mutations in Twinkle primase-helicase cause Perrault syndrome with neurologic features. Neurology. 2014;83(22):2054–61.

10. Chatzispyrou IA, Alders M, Guerrero-Castillo S, Zapata Perez R, Haagmans MA, Mouchiroud L, et al. A homozygous missense mutation in ERAL1, encoding a mitochondrial rRNA chaperone, causes Perrault syndrome. Hum Mol Genet. 2017. Epub 2017/04/28.

11. Demain LA, Urquhart JE, O’Sullivan J, Williams SG, Bhaskar SS, Jenkinson EM, et al. Expanding the genotypic spectrum of Perrault syndrome. Clinical genetics. 2017;91(2):302–12. Epub 2016/03/13.

12. Lerat J, Jonard L, Loundon N, Christin-Maitre S, Lacombe D, Goizet C, et al. An Application of NGS for Molecular Investigations in Perrault Syndrome: Study of 14 Families and Review of the Literature. Hum Mutat. 2016;37(12):1354–62. Epub 2016/09/22.

13. Ferdinandusse S, Denis S, Mooyer PA, Dekker C, Duran M, Soorani-Lunsing RJ, et al. Clinical and biochemical spectrum of D-bifunctional protein deficiency. Ann Neurol. 2006;59(1):92–104. Epub 2005/11/10.

14. McMillan HJ, Worthylake T, Schwartzentruber J, Gottlieb CC, Lawrence SE, MacKenzie A, et al. Specific combination of compound heterozygous mutations in 17β-hydroxysteroid dehydrogenase type 4 (HSD17B4) defines a new subtype of D-bifunctional protein deficiency. Orphanet J Rare Dis. 2012;7:90.

15. Holzmann J, Frank P, Loffler E, Bennett KL, Gerner C, Rossmanith W. RNase P without RNA: identification and functional reconstitution of the human mitochondrial tRNA processing enzyme. Cell. 2008;135(3):462–74. Epub 2008/11/06.

16. Vilardo E, Nachbagauer C, Buzet A, Taschner A, Holzmann J, Rossmanith W. A subcomplex of human mitochondrial RNase P is a bifunctional methyltransferase-extensive moonlighting in mitochondrial tRNA biogenesis. Nucleic Acids Res. 2012;40(22):11583–93.

17. Banka S, Blom HJ, Walter J, Aziz M, Urquhart J, Clouthier CM, et al. Identification and characterization of an inborn error of metabolism caused by dihydrofolate reductase deficiency. Am J Hum Genet. 2011;88(2):216–25.

18. Carr IM, Flintoff KJ, Taylor GR, Markham AF, Bonthron DT. Interactive visual analysis of SNP data for rapid autozygosity mapping in consanguineous families. Hum Mutat. 2006;27(10):1041–6.

19. NHLBI GO Exome Sequencing Project (ESP). Exome Variant Server. Seattle, WA [cited 2015 09/07]; Available from: http://evs.gs.washington.edu/EVS/.

20. Sherry ST, Ward MH, Kholodov M, Baker J, Phan L, Smigielski EM, et al. dbSNP: the NCBI database of genetic variation. Nucleic Acids Res. 2001;29(1):308–11.

21. Action on Hearing Loss. Levels of hearing loss. Action on hearing loss; 2017 [cited 2017 12/06]; Available from: https://www.actiononhearingloss.org.uk/your-hearing/about-deafness-and-hearing-loss/glossary/levels-of-hearing-loss.aspx.

22. Aiman J, Smentek C. Premature ovarian failure. Obstet Gynecol. 1985;66(1):9–14. Epub 1985/07/01.

23. Kumar P, Henikoff S, Ng PC. Predicting the effects of coding non-synonymous variants on protein function using the SIFT algorithm. Nat Protoc. 2009;4(7):1073–81.

24. Adzhubei IA, Schmidt S, Peshkin L, Ramensky VE, Gerasimova A, Bork P, et al. A method and server for predicting damaging missense mutations. Nat Methods. 2010;7(4):248–9. Epub 2010/04/01.

25. Schwarz JM, Cooper DN, Schuelke M, Seelow D. MutationTaster2: mutation prediction for the deep-sequencing age. Nat Methods. 2014;11(4):361–2. Epub 2014/04/01.

26. Exome Aggregation Consortium (ExAC). Exome Aggregation Consortium (ExAC). Cambridge, MA [cited 2015 08/07]; Available from: http://exac.broadinstitute.org.

27. Narasimhan VM, Hunt KA, Mason D, Baker CL, Karczewski KJ, Barnes MR, et al. Health and population effects of rare gene knockouts in adult humans with related parents. Science. 2016;352(6284):474–77.

28. Deutschmann AJ, Amberger A, Zavadil C, Steinbeisser H, Mayr JA, Feichtinger RG, et al. Mutation or knock-down of 17beta-hydroxysteroid dehydrogenase type 10 cause loss of MRPP1 and impaired processing of mitochondrial heavy strand transcripts. Hum Mol Genet. 2014;23(13):3618–28. Epub 2014/02/20.

29. Howard MJ, Lim WH, Fierke CA, Koutmos M. Mitochondrial ribonuclease P structure provides insight into the evolution of catalytic strategies for precursor-tRNA 5’ processing. Proc Natl Acad Sci U S A. 2012;109(40):16149–54. Epub 2012/09/20.

30. Reinhard L, Sridhara S, Hallberg BM. Structure of the nuclease subunit of human mitochondrial RNase P. Nucleic Acids Res. 2015;43(11):5664–72. Epub 2015/05/09.

31. Rossmanith W. Processing of human mitochondrial tRNA(Ser( AGY))(GCU): A novel pathway in tRNA biosynthesis. J Mol Biol. 1997;265(4):365–71.

32. Ehret G. Development of absolute auditory thresholds in the house mouse (Mus musculus). Journal of the American Audiology Society. 1976;1(5):179–84. Epub 1976/03/01.

33. Sendin G, Bulankina AV, Riedel D, Moser T. Maturation of ribbon synapses in hair cells is driven by thyroid hormone. The Journal of neuroscience: the official journal of the Society for Neuroscience. 2007;27(12):3163–73. Epub 2007/03/23.

34. Lesus J, Perkins G, Lysakowski A. Structural Analysis of Inner Ear Hair Cell Mitochondria Near the Striated Organelle. The FASEB Journal. 2017;31(1 Supplement):740.18-.18.

35. Lysakowski A, Goldberg JM. A regional ultrastructural analysis of the cellular and synaptic architecture in the chinchilla cristae ampullares. The Journal of comparative neurology. 1997;389(3):419–43. Epub 1997/12/31.

36. Brzezniak LK, Bijata M, Szczesny RJ, Stepien PP. Involvement of human ELAC2 gene product in 3’ end processing of mitochondrial tRNAs. RNA biology. 2011;8(4):616–26. Epub 2011/05/20.

37. Rossmanith W. Localization of human RNase Z isoforms: dual nuclear/mitochondrial targeting of the ELAC2 gene product by alternative translation initiation. PloS one. 2011;6(4):e19152. Epub 2011/05/12.

38. Ojala D, Montoya J, Attardi G. tRNA punctuation model of RNA processing in human mitochondria. Nature. 1981;290(5806):470–4. Epub 1981/04/09.

39. Rossmanith W. Of P and Z: mitochondrial tRNA processing enzymes. Biochim Biophys Acta. 2012;1819(9-10):1017–26.

40. Zschocke J. HSD10 disease: clinical consequences of mutations in the HSD17B10 gene. JIMD. 2012;35(1):81–9.

41. Metodiev MD, Thompson K, Alston CL, Morris AA, He L, Assouline Z, et al. Recessive Mutations in TRMT10C Cause Defects in Mitochondrial RNA Processing and Multiple Respiratory Chain Deficiencies. Am J Hum Genet. 2016;98(5):993–1000. Epub 2016/05/03.

42. Chatfield KC, Coughlin CR, Friederich MW, Gallagher RC, Hesselberth JR, Lovell MA, et al. Mitochondrial energy failure in HSD10 disease is due to defective mtDNA transcript processing. Mitochondrion. 2015;21:1–10.

43. Blomen VA, Majek P, Jae LT, Bigenzahn JW, Nieuwenhuis J, Staring J, et al. Gene essentiality and synthetic lethality in haploid human cells. Science. 2015;350(6264):1092–6. Epub 2015/10/17.

44. Wang T, Birsoy K, Hughes NW, Krupczak KM, Post Y, Wei JJ, et al. Identification and characterization of essential genes in the human genome. Science. 2015;350(6264):1096–101. Epub 2015/10/17.

45. Rauschenberger K, Schöler K, Sass JO, Sauer S, Djuric Z, Rumig C, et al. A non - enzymatic function of 17*β*-hydroxysteroid dehydrogenase type 10 is required for mitochondrial integrity and cell survival. EMBO Mol Med. 2010;2(2):51–62.

46. Rackham O, Busch JD, Matic S, Siira SJ, Kuznetsova I, Atanassov I, et al. Hierarchical RNA Processing Is Required for Mitochondrial Ribosome Assembly. Cell reports. 2016;16(7):1874–90. Epub 2016/08/09.

47. Vilardo E, Rossmanith W. Molecular insights into HSD10 disease: impact of SDR5C1 mutations on the human mitochondrial RNase P complex. Nucleic Acids Res. 2015;43(10):5112–9. Epub 2015/06/21.

48. Riley LG, Rudinger-Thirion J, Schmitz-Abe K, Thorburn DR, Davis RL, Teo J, et al. LARS2 Variants Associated with Hydrops, Lactic Acidosis, Sideroblastic Anemia, and Multisystem Failure. JIMD Reports. 2015;24:1–9.

49. Nishio SY, Takumi Y, Usami SI. Laser-capture micro dissection combined with next-generation sequencing analysis of cell type-specific deafness gene expression in the mouse cochlea. Hearing research. 2017;348:87–97. Epub 2017/03/07.

50. Smith MJ, Beetz C, Williams SG, Bhaskar SS, O’Sullivan J, Anderson B, et al. Germline mutations in SUFU cause Gorlin syndrome-associated childhood medulloblastoma and redefine the risk associated with PTCH1 mutations. J Clin Oncol. 2014;32(36):4155–61.

51. Kunkel TA, Roberts JD, Zakour RA. Rapid and efficient site-specific mutagenesis without phenotypic selection. Methods Enzymol. 1987;154:367–82.

52. O’Keefe RT, Norman C, Newman AJ. The Invariant U5 snRNA Loop 1 Sequence Is Dispensable for the First Catalytic Step of pre-mRNA Splicing in Yeast. Cell. 1996;86(4):679–89.

53. Rossmanith W, Tullo A, Potuschak T, Karwan R, Sbisa E. Human mitochondrial tRNA processing. J Biol Chem. 1995;270(21):12885–91.

